# Acceleration of Nucleotide Semi-Global Alignment with Adaptive Banded Dynamic Programming

**DOI:** 10.1101/130633

**Authors:** Hajime Suzuki, Masahiro Kasahara

## Abstract

**Motivation:** Pairwise alignment of nucleotide sequences has previously been carried out using the seed-and-extend strategy, where we enumerate seeds (shared patterns) between sequences and then extend the seeds by Smith-Waterman-like semi-global dynamic programming to obtain full pairwise alignments. With the advent of massively parallel short read sequencers, algorithms and data structures for efficiently finding seeds have been extensively explored. However, recent advances in single-molecule sequencing technologies have enabled us to obtain millions of reads, each of which is orders of magnitude longer than those output by the short-read sequencers, demanding a faster algorithm for the extension step that accounts for most of the computation time required for pairwise local alignment. Our goal is to design a faster extension algorithm suitable for single-molecule sequencers with high sequencing error rates (e.g., 10-15%) and with more frequent insertions and deletions than substitutions.

**Results:** We propose an adaptive banded dynamic programming algorithm for calculating pairwise semi-global alignment of nucleotide sequences that allows a relatively high insertion or deletion rate while keeping band width relatively low (e.g., 32 or 64 cells) regardless of sequence lengths. Our new algorithm eliminated mutual dependences between elements in a vector, allowing an efficient Single-Instruction-Multiple-Data parallelization. We experimentally demonstrate that our algorithm runs approximately 5× faster than the extension alignment algorithm in NCBI BLAST+ while retaining similar sensitivity (recall).

We also show that our extension algorithm is more sensitive than the extension alignment routine in DALIGNER, while the computation time is comparable.

**Availability:** The implementation of the algorithm and the benchmarking scripts are available at https://github.com/ocxtal/adaptivebandbench.

## 1 Introduction

In the past decade, technological improvement in the DNA sequencing field has been remarkable. Single-molecule sequencers, often called third-generation sequencers, achieved more than tenfold improvement in their read lengths. The commercially available third-generation sequencers, such as the PacBio Sequel and Oxford Nanopore MinION, can yield reads of 20 kb or even longer (Gordon *et al.* (2016) and Jain *et al.* (2017)), whereas the longest practical read lengths of the Sanger sequencers were around 1 kb. This creates the need to process huge numbers of reads longer than 20 kb, but algorithms for such purposes were not well developed before third-generation sequencers became common.

Most genome analyses using massively parallel sequencers start with aligning reads against themselves (for *de novo* assembly in whole genome re-sequencing research; Chin *et al.* (2013), Koren *et al.* (2017)) or reference sequences (for other reference-guided analyses, e.g. exome sequencing and RNA-seq; Pabinger *et al.* (2014), Ozsolak and Milos (2011)). Therefore, faster algorithms for pairwise alignment are crucial in order to accelerate most types of genomic analysis. However, it has recently been shown that the near-quadratic time bounds for computing edit distance cannot be improved, and are expected to be a limiting factor (unless the strong exponential time hypothesis is false; Backurs and Indyk (2015)). Therefore, it is desirable to design fast heuristic algorithms for pairwise alignment. Typical heuristic algorithms for pairwise local alignment first find short exact matches called “seeds,” then extend them using a semi-global variant of pairwise local alignment algorithms, such as the Smith-Waterman-Gotoh algorithm (SWG; Smith and Waterman (1981); Gotoh (1982)). This idea, the seed-and-extend strategy, was employed in the classical Basic Local Alignment Search Tool (BLAST; Altschul *et al.* (1990)), and more recently in long-read alignment programs for third-generation sequencers, such as BWA-MEM (Li, 2013), BLASR (Chaisson and Tesler, 2012), DALIGNER (Myers, 2014), and GraphMap (Sović *et al.*, 2016).

In the era of second-generation sequencers, researchers focused on the development of fast algorithms for identifying seed matches because the seed-matching stage was the most time-consuming step. Approximate string-matching algorithms based on Baeza-Yates algorithm (Navarro and Ricardo, 1998) were used in the early days (MAQ (Li *et al.*, 2008) and BFAST (Homer *et al.*, 2009)), and later an exact substring-matching algorithm based on Burrows-Wheeler transform (Burrows and Wheeler, 1994) and an auxiliary data structure proposed by Ferragina and Manzini (2000) were adopted by many sequence alignment programs, such as BWA (Li and Durbin, 2009) and Bowtie 2 (Langmead and Salzberg, 2012).

However, as read length increases, the extension step accounts for a greater proportion of computation time for pairwise alignment, and therefore a faster extension algorithm is required. Commonly used techniques for accelerating the extension alignment calculate only the values of cells in a small region of the Dynamic Programming (DP) matrix in the SWG algorithm; we first create a heuristic estimate of a region through which the optimal path of the pairwise alignment may travel, and compute only the values of the cells in that region. As the region gets smaller, computation time for the extension step decreases, but we run a higher risk of missing an optimal path (alignment). This technique was first proposed by Chao *et al.* (1992), and later adopted by many local alignment programs such as BWA-MEM (Li, 2013) or BLASR (Chaisson and Tesler, 2012) as “banded DP.” Nonetheless, the required band width for this approach is still too large when an outermost seed is far from the end of a sequence; longer a region without seeds become, more the optimal alignment path drifts off the diagonal.

To avoid this problem, BLASR (Chaisson and Tesler, 2012) and minimap2 (Li, 2017) first chains seeds in order to estimate where the optimal path travels through in the DP matrix, and then do static banded DP to calculate detailed alignments. This approach should work well if sufficiently long and correct seeds are identified uniformly across sequences in the seeding step, but in reality there might not be the case: (1) we need to find seeds even in duplicated or repetitive regions when there is a large chunk of such sequences in target sequences, although such regions are usually masked implicitly or explicitly by alignment programs. (2) we need to use shorter and sensitive seeds under the presence of abundant indel errors, which increases the computation time. An extension algorithm that works well in such situations is demanded.

Another heuristic for the extension step was introduced in BLAST, which we denote by the “BLAST X-drop DP algorithm.” The BLAST X-drop DP algorithm continues to extend alignments until all cells in the forefront have a score less than the current maximum minus *X* by implicitly assuming that valid alignments do not include parts of alignments with scores that have values under –*X*. The algorithm successfully reduces the region in the DP matrix to be calculated when *X* is sufficiently small, but there was no parallel implementation for the algorithm.

Previous approaches for accelerating the SWG algorithm by a constant factor also include the use of Single-Instruction-Multiple-Data (SIMD) operations that increase the number of cells processed per unit operation. Examples of such approaches include Wozniak’s (Wozniak, 1997), Rognes’s (Rognes and Seeberg, 2000), and Farrar’s (Farrar, 2007) methods. Farrar’s striped vectorization (parallelization) successfully accelerated the calculation of a rectangular DP matrix of the SWG algorithm, and has been adopted by many local alignment programs (e.g., MOSAIK2 (Lee *et al.*, 2014), BWA (Li and Durbin, 2009) and Bowtie 2 (Langmead and Salzberg, 2012)) and libraries (e.g., SSW library (Zhao *et al.*, 2013), Parasail (Daily, 2016)).

Here, we propose a new adaptive banded DP algorithm wherein the shape of the band is determined dynamically during extension and the band width is fixed regardless of sequence lengths (Fig 1), and therefore the computation time is linear to the sequence length, while the loss in sensitivity is small. Adaptive banded DP uses SIMD instructions on general-purpose processors in a novel way, with anti-diagonal forefront vector placement, and supports an X-drop-like heuristic to identify the ends of alignments. We demonstrate that adaptive banded DP is the fastest affine gap penalty semi-global alignment algorithm for extension alignment for long single-molecule sequencing reads.

**Fig. 1.**
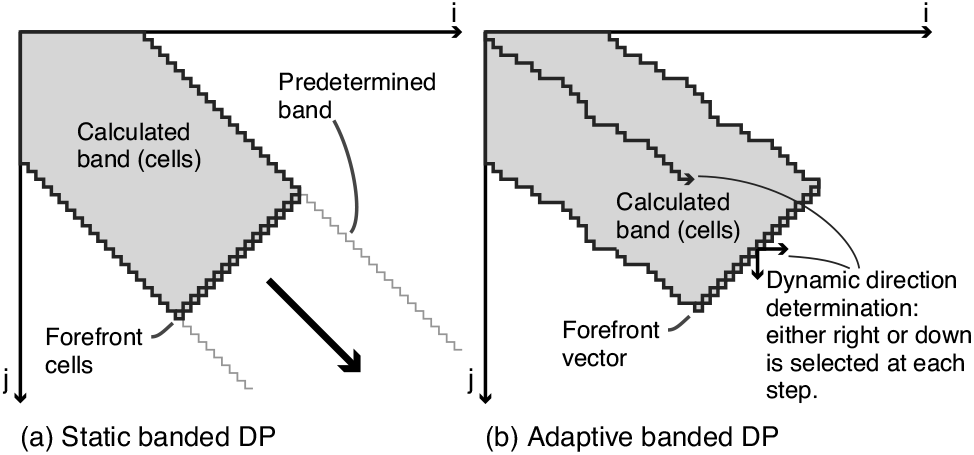
Comparison of Static and Adaptive Banded Dynamic Programming Algorithms. (a) Illustration of the static banded DP algorithm: The band is determined in advance of the matrix calculation. The cells at the forefront of the band are placed in the antidiagonal direction in this example, whereas common implementations, for example, the extension DP routines in BWA-MEM and BLASR, adopt horizontal or vertical forefront cell compositions. (b) Illustration of the adaptive banded DP algorithm: The band is determined dynamically during extension, in contrast to the static banded DP algorithm.

## 2 Methods

### 2.1 Semi-global alignment of nucleotide sequences

First, let us define the nucleotide semi-global alignment problem. Let *a* = *a*_0_*a*_1_…*a*_|*a*|−1_ and *b* = *b*_0_*b*_1_…*b*_|*b*| − 1_ be strings over an alphabet Σ = {A, C, G, T}. The problem formulation is to calculate a coordinate (*n*, *m*) and a corresponding “alignment,” or an edit path from (0, 0) to (*n,m*) that consists of {match, insertion, deletion}, that maximizes the sum of substitution scores and gap penalties. The substitution scores are defined over a pair of letters: *s*(*p*, *q*) where *p,q* ∈ Σ (called “score matrix”), and the gap penalty function is expressed in an integer linear form: *g*(*k*) = *G*_*o*_ + *k* · *G*_*e*_, where *G*_*o*_ ≥ 0, *G*_*e*_ > 0 and *k* is the length of contiguous gaps (called the “affine gap penalty function”). The problem appears as a subproblem in the extension stage of the seed-and-extend algorithm.

This formulation of the semi-global alignment problem is usually solved with a variant of the Smith-Waterman-Gotoh (Smith and Waterman, 1981; Gotoh, 1982) algorithm, where the initial values (the scores at the top and left edges in the DP matrix) are modified to the gap penalties from the origin. This modification fixes the starting cell of resulting alignments to the origin of the matrix. The ends of alignments remain open (not fixed), as in the original SWG algorithm; it initiates traceback from the cell with the highest score. We use a general 4 × 4 score matrix throughout the paper unless otherwise specified. However, the match-mismatch scoring model (wherein a score matrix is characterized by a pair of integers, (*M*, *X*), where *M* is a match score and *X* is a penalty score) is a special case of the general score matrix, and therefore most discussions hereafter also hold true for the match-mismatch scoring model. The recurrence relations of the DP matrices used in the SWG algorithm are shown in Equation 1, where *S* is a score matrix and *E* and *F* are used to calculate gap penalties in the horizontal and vertical directions, respectively.

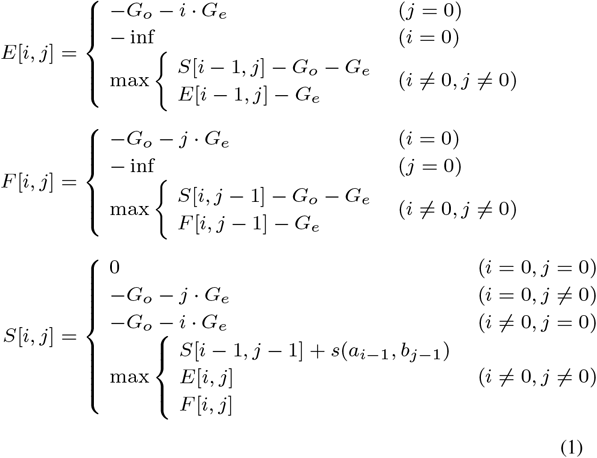

We use as — inf a sufficiently small value within the range of the integer type used in an actual implementation. Note that, while this formulation of the SWG algorithm might differ from those that appear in previous papers, it is mathematically equivalent to the corrected version of the original SWG algorithm (Flouri *et al.*, 2015).

### 2.2 Adaptive banded DP

In 1992, Chao *et.al.* proposed a method called “banded DP” as an acceleration technique for the global alignment DP algorithms. This method reduced the time and memory necessary for computation by narrowing the region of the DP matrix to be calculated. The idea was later simplified to fill a pre-determined, diagonally placed rectangular band with a constant width, which we hereafter call “static banded DP.” The algorithm was applied to many semi-global alignment implementations, such as SeqAn (Döring *et al.*, 2008), Parasail library (Daily, 2016), the refinement alignment in the SSW library (Zhao *et al.*, 2013), and the extension alignment in BWA-MEM (Li, 2013) and BLASR (Chaisson and Tesler, 2012). A vector-oriented parallelization of static banded DP was proposed by Kimura *et al.* (2012) for calculating the edit-distance. They adopted anti-diagonally placed vectors with a constant width (e.g., 64 cells) to calculate the cells in a vector simultaneously.

A new adaptive banded DP algorithm that we propose here adopts a similar band-narrowing approach. However, in contrast to static banded DP, where the narrowed region is determined statically (i.e., before filling cells in the DP matrix), our algorithm determines the narrowed region dynamically as we calculate cells in the DP matrix. A *forefront vector* is a set of cells placed along with the anti-diagonal direction (Fig 1b). The values of the cells in a forefront vector are calculated in parallel using SIMD operations. The width of the forefront vector is a constant, which is ideally the width of SIMD registers but may also be multiples of it. Other width is also possible, albeit less efficient. The forefront vector is initially at the origin of the DP matrix. It advances either rightward or downward iteratively until it reaches the end of alignment. Figure 3 provides a more detailed overview of our algorithm. The banded region with a constant width (the number of anti-diagonally aligned cells; denoted as W) is created by iteratively pushing the forefront DP vectors towards the diagonal direction (i.e., rightward or downward). The three forefront DP vectors, *S*_*V*_, *E*_*V*_, and *F*_*V*_, hold the cells in an anti-diagonal line (Fig 3(a)). At each step, the three forefront DP vectors move either rightward or downward. On every step, the advancing direction of the forefront vectors is determined by comparing the two edge cells (*S*_*V*_ [0] and *S*_*V*_ [*W* − 1]) to ensure that the difference between the two edge cells in the next forefront *S*_*V*_ vector is smaller. In other words, the next vectors are placed rightward when *S*_*V*_ [0] ≥ *S*_*V*_ [*W* − 1] and downward when *S*_*V*_ [0] < *S*_*V*_ [*W* − 1] (Fig 3(b, c)). When the forefront vectors move, the *E*_*V*_, *F*_*V*_, or *S*_*Y*_ are updated according to the formula (Fig 3(d)). First, the new *E*_*V*_ and *F*_*V*_ vectors are derived from the previous *E*_*V*_, *F*_*V*_, and *S*_*V*_ vectors; the new *S*_*V*_ vector is then calculated from the current *E*_*V*_ and *F*_*V*_ vectors, the second-previous *S*_*V*_ vector, and a score profile vector (an array of substitution scores). Since there are no dependences between the cells in the forefront vectors in each operation, the update procedure can be implemented with Single-Instruction-Multiple-Data (SIMD) instructions, keeping the vectors on SIMD registers.

**Fig. 2.**
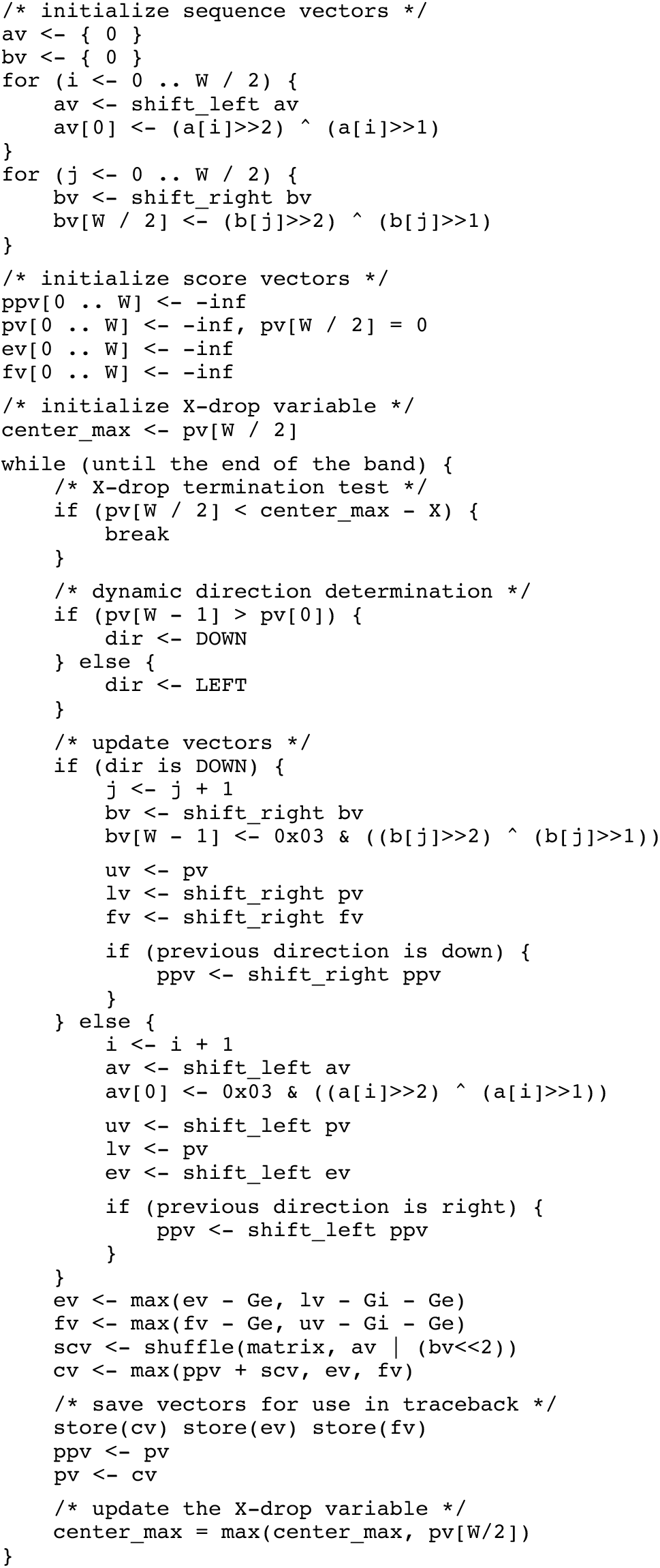
Pseudocode of the Adaptive Banded DP Algorithm. ppv, pv, ev, and fv represent the second-previous *S*_*V*_ vector, the previous *S*_*V*_ vector, the *E*_*V*_ vector, and the *F*_*V*_ vector, respectively. The two subsequent vectors are denoted as av and bv. The X-drop threshold is denoted as *X*. The three binary operators on vectors − +, –, and max - are element-wise addition, subtraction, and maximum, respectively. The shift_left and shift_right operators shift elements in a vector leftward and rightward, respectively, by one column. The shuffle operation takes an element vector as the first argument and an index vector as the second argument. The ≪ operator in the shuffle represents the bitwise leftward shift of each element in the vector.

**Fig. 3.**
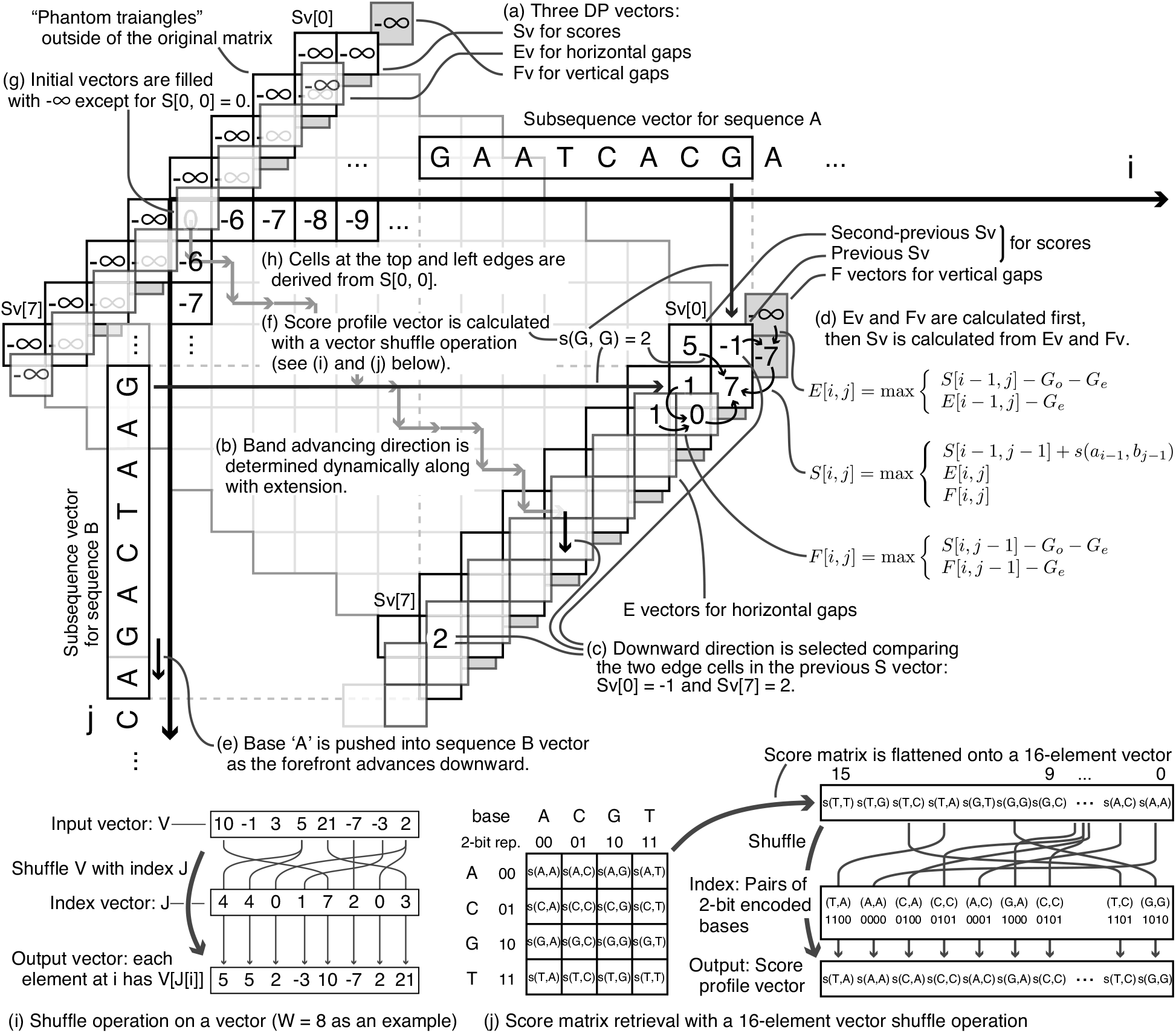
Overview of the adaptive banded DP algorithm, where *W* = 8. (a) Vector placement: Three DP vectors (*S*_*V*_, *E*_*V*_, and *F*_*V*_) are placed in the anti-diagonal direction of the DP matrix, holding the corresponding parts of the original *S*, *E*, and *F* matrices. (b, c) Determining the advancing direction of the band: The advancing direction of the band is determined on every vector update by comparing the two edge cells in the *S*_*V*_ vector. (d) Vector update procedure: The vector update procedure consists of three update operations, each of which corresponds to the update formula of the original semi-global alignment algorithm (Eq 1). (e) Subsequence vectors: Two subsequence vectors are placed on the top and left sides of the matrix. They move according to the advancing direction of the band. (f, i, j) Score profile vector calculation: The score profile vector, an array of *s* (·, ·) values, is calculated using a vector shuffle instruction (i), which is an indexed element retrieval on a vector. The index vector and the input element vector are respectively composed of pairs of 2-bit encoded bases, and the flattened 4 × 4 substitution matrix (j). (g, h) Initialization of vectors: The three, *S*_*V*_, *E*_*V*_, and *F*_*V*_ vectors and the second-previous *S*_*V*_ vector are initialized with – inf except for S [0, 0] = 0. This setting results in proper initial values aligning on the top and leftmost lines.

In our algorithm, the score profile vector is also generated in a SIMD-vectorized manner using a vector shuffle operation. The vector shuffle operation can retrieve multiple elements (e.g., 16 elements) from a given array of a fixed size (e.g., 16) in a single operation; it does a simple table-lookup from the 16-element array multiple times (e.g., 16 times) in parallel (Fig 3(i)). We concatenate the pair of 2-bit encoded bases into one value ranging from 0 to 15, and use the array of these values as the index vector and a flattened 4 × 4 score matrix as the input vector of the shuffle operation (Fig 3(f, j)). To generate the index vector efficiently, we retain two subsequences of length *W* on vector registers. Each base is represented in a 2-bit binary code, where {A, C, G, T} are mapped to {00,01,10,11}, respectively. Every time the forefront vector moves, either of the two sequence vectors is shifted left or right by one according to the advancing direction (Fig 3(e)). The conversion from an ASCII code to the 2-bit encoded binary is performed when a base comes into the sequence vector; the conversion is performed based on a simple and well-known formula 0×03 · ((*c* ≫ 2) ⊕ (*c* ≫ 1)) where c is a 8-bit ASCII-encoded character and ·, ⊕, and ≫ are respectively bitwise AND, bitwise XOR, and logical shift operations. To our knowledge, the technique has not been reported in the literature, nor could we find the origin of the technique. Note that the conversion works correctly regardless of the case of an input character, since the output 2-bit pattern depends on only a subset of input bits whose uppercase character and corresponding lowercase character are identical.

The head of the band, the top-left triangular corner of the matrix, is handled in a special way. We added two phantom triangular regions to reshape the corner in order to maintain a constant width. The initial vectors are placed at the top-left edge of the augmented band, whose center cell^1^ is aligned to the cell at the origin (0, 0). The initial values in the vectors are set to — inf, or a sufficiently small value within the range of the cell variable, except that the cell at (0, 0) is set to 0 in order to derive proper initial values in the first column (*i* = 0) and the first row (*j* = 0) in the DP matrix. Figure 3(f) shows an example of the initial and derived values with the score parameters (*M*, *X*, *G*_*o*_, *G*_*e*_) = (2, 3, 5, 1).

Finally, we describe a heuristic to terminate the band extension, which is similar to an algorithm introduced in the BLAST X-drop DP algorithm. To avoid unfruitful extension through an unmatched region beyond the true end of the matches, the BLAST (Altschul *et al.*, 1990) and subsequent programs such as BWA-MEM (Li, 2013) and LAST (Kielbasa *et al.*, 2011) adopted a heuristic algorithm called X-drop termination in their semiglobal alignment DP routines. This algorithm terminates the extension when all of the forefront cell scores are smaller than the current maximum score by at least *X*. Since there is no way to calculate the maximum values in a vector efficiently without time-consuming folding with [log *W*] steps, we adopted a slightly modified version of the X-drop heuristic to avoid inefficiency. Our algorithm does not find the maximum value in the forefront vector as the original X-drop heuristic does. Instead, the extension is terminated when the score of the center cell at the forefront of the band becomes smaller than the current maximum score in the previously calculated center cell by at least *X*.

The whole adaptive banded DP algorithm is shown in pseudocode in Figure 2, where `ppv, pv, ev,` and `fv` correspond to the second-previous *S*_*V*_ vector, and previous *S*_*V*_, *E*_*V*_, and *F*_*V*_ vectors in Figure 3, respectively. The two subsequent vectors are denoted as `av` and `bv`.

### 2.3 Relation to existing algorithms

Our approach, which uses the anti-diagonal vector for parallelization, is similar to the parallel SWG algorithm with SIMD instructions by Wozniak (1997). Wozniak’s algorithm, which targeted mainly protein sequences, had a bottleneck in its serial (unparallelized) lookups of a score matrix; it was later superseded by more efficient parallel algorithms by Rognes and Seeberg (2000) and Farrar (2007). In Rognes’s algorithm and Farrar’s algorithm, the DP vectors are vertical (or horizontal) in the DP matrix so that the score profile vector can be calculated by choosing (loading) one of the precalculated four (or 20) score profile vectors that correspond to the four nucleotides (20 amino acids). The precalculated score profile vectors were effective in eliminating the serial score-matrix lookups in Wozniak’s algorithm, but entailed precalculation and additional memory consumption for the vectors.

The score profile vector calculation with an SIMD shuffle operation was first proposed by Wang *et al.* (2014) in their XSW program, which adopted Farrar’s SIMD-vectorized SWG algorithm. They use a pair of 16-element vector shuffle operations to calculate a 16-element score vector from a single row of 26 × 26 score matrix in an on-the-fly manner. The parallel score vector calculation in our algorithm can be considered a further extension of Wang’s approach; it accepts arbitrary base pairs as the input and eliminates the indexed memory access required in Wang’s algorithm when fetching a row from the score matrix. We noticed that the 4 × 4 score matrix for nucleotide alignment fits perfectly in a single 16-element vector, while the score matrix for protein alignment does not. This enabled us to combine the anti-diagonal-parallel approach and the vectorized score profile calculation in an efficient way.

We also found that a hardware-based semi-global alignment algorithm proposed in a patent by McMillen and Ruehle (2015) also adopted a constant-width band that moves dynamically according to the values in calculated cells. However, further details about how to move the forefront vector are not described even in the mode of operation of invention. We speculate from the patent document that their method moves the forefront vector so that the cell with the maximum value comes closer to the center; thus, our approach likely differs in this respect.

## 3 Results

We implemented our adaptive banded DP algorithm for x86_64 processors with Streaming SIMD Extension 4.1 (SSE4.1) instruction sets. We used the 16-bit-wide variables; eight values were retained in a single xmm SIMD register and processed simultaneously during the extension. The band width *W* was set to multiples of 8 and determined at the compile time by passing the constant value as a macro definition to the compiler. All benchmarking programs were implemented in the C programming language and compiled with gcc-5.3.1 and executed on a cluster node with dual Intel Xeon X5650 (Westmere-EP) 2.67 GHz processors.

### 3.1 Recall benchmarks on simulated reads

Since our algorithm calculates only the cells in the narrowed regions, it may miss the optimal alignment that could be identified by the original semiglobal DP algorithm that calculates the full DP matrix. To establish that the sensitivity of our algorithm is similar to that of the original semi-global DP algorithm, we used a simulation to compare the optimal score with the corresponding alignments identified by our algorithm and the original semi-global DP algorithm under several conditions. In the simulation, we generated simulated reads from a reference genome. We then aligned the simulated reads against the reference genome using both algorithms, mimicking atypical scenario in resequencing studies. We assumed that two parameters, the band width *W* and the score parameters, had strong effects on the accuracy and sensitivity of alignment. Throughout the experiments, we used the simple match-mismatch score model, which is represented as a tuple of four non-negative integers (*M*, *X*, *G*_*o*_, *G*_*e*_), where *M* and *X* are a match reward and a mismatch penalty, and *G*_*o*_ and *G*_*e*_ are the coefficients of the affine-gap penalty function. Recall that the match-mismatch score model is a special case of the 4 × 4 score matrix model as described in the Methods section. The X-drop threshold was set to 40 in the benchmarks on simulated reads. Sets of 1,000 sequence pairs were generated from the *Escherichia coli* reference genome (accession No.: NC000913) with PBSIM, a PacBio long-read simulator (Ono *et al.*, 2013). Each pair consisted of a simulated PacBio read and the corresponding region in the reference genome, to each of which a random sequence of 200 bp was appended in order to mimic a semi-global alignment scenario. In experiments that followed, we characterized a set of sequences by mean read length *L* and mean identity *I*. The maximum, minimum, and standard deviation of these two parameters were set to 1.05*L*, 0.95*L*, and 0.05*L*, and *I* + 0.01, *I* − 0.01, and 0.01, respectively. The statistics of one of the sets of simulated reads are provided as an example in Table (a) in Figure 4.

**Fig. 4.**
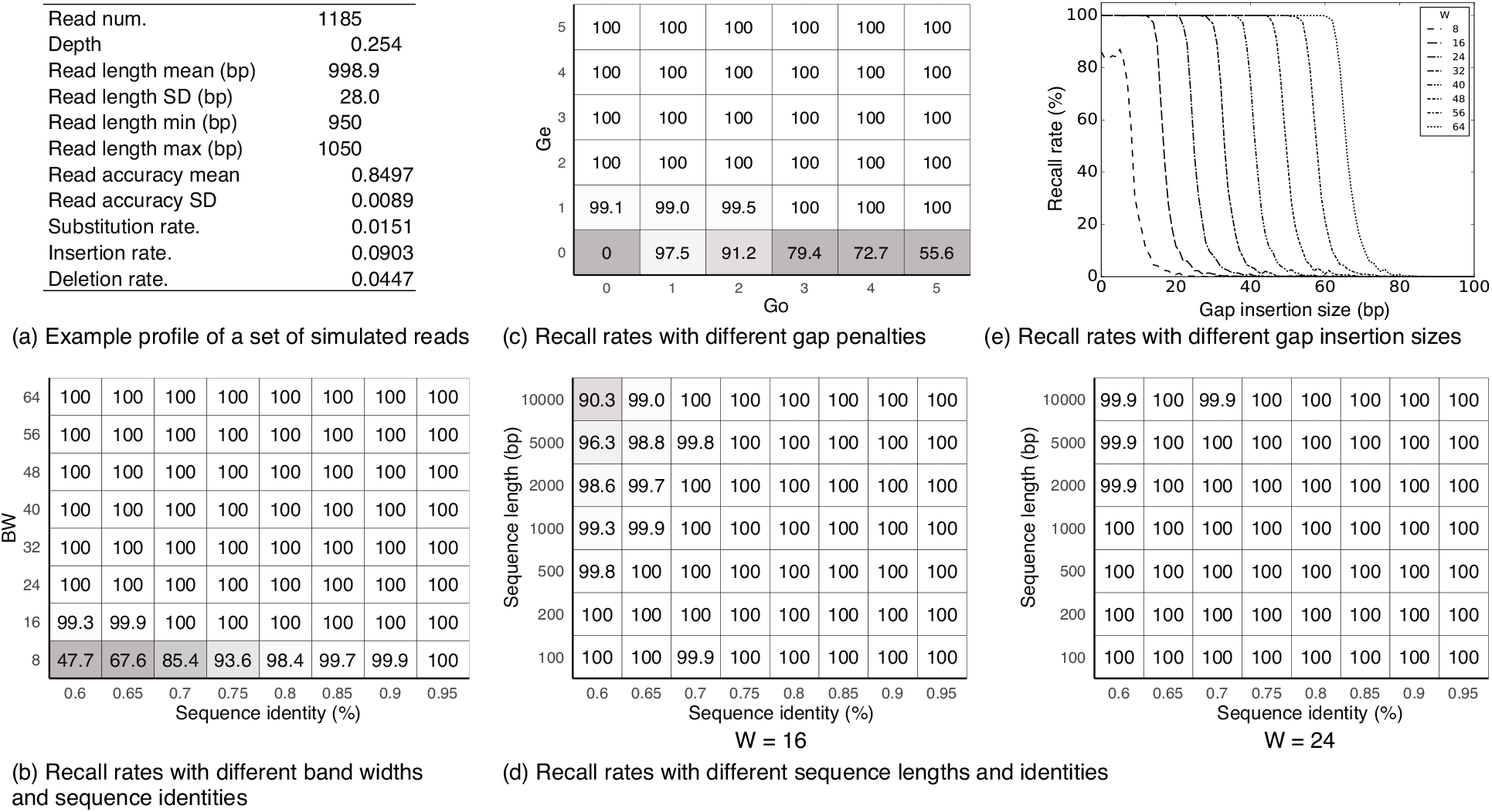
Results of the recall benchmarks (a) Statistical profiles of one sample of the simulated read sets: The length and the identity were set to 1,000 bp and 0.85, with their (SD, max, min) set to (50, 1050, 950) and (0.01, 0.86, 0.84), respectively. (b) Recall rate (in percent) with different band widths and sequence identities: The mean sequence length L and score parameters were set to 1, 000 and (1, 2, 2, 1), respectively. 100.0 is displayed as 100. (c) Recall rate (in percent) with different gap penalties: The match award and mismatch penalty were fixed at (*M,X*) = (2, 2). The mean sequence length and identity were set to 1, 000 and 0.75, respectively. 100.0 is displayed as 100.(d) Recall rate (in percent) with different sequence lengths and identities with two band widths: The result of *W* = 16 is shown on the left and *W* = 24 on the right. The scoring parameters were set to (1, 2, 2, 1). 100.0 is displayed as 100. (e) Recall rate (in percent) with different gap insertion sizes: The mean sequence length and identity were set to 1,000 and 0.75, respectively. A single contiguous gap with a length within [0, 100] was inserted into the reference-side sequence at a position within [100, 600] from its head. The scoring parameters were set to (*M,X,G*_*o*_, *G*_*e*_) = (1, 2, 2, 1).

#### 3.1.1 Effect of the band width on the recall rate

We first evaluated the effect of the band width *W* on the recall rate of the algorithm. Sequence pairs with various identities within the range of [0.6, 0.95] were aligned using the adaptive banded DP algorithm with band widths ranging from 8 to 64 with a step of 8. The mean sequence length and the scoring parameters were set to *L* = 1, 000 and (*M*, *X*, *G*_*o*_, *G*_*e*_) = (1, 2, 2, 1), respectively. The result is shown in Figure 4(b), demonstrating that our algorithm can perfectly identify the optimal scores and paths when the band width is greater than or equal to 24 and when the sequence identity is 0.6 or greater. The results are still nearly perfect when *W* = 16, while the recall rate drops significantly with *W* = 8 and sequence identities lower than 0.75.

#### 3.1.2 Effect of gap penalties on recall rate

The algorithm examines only the scores of the edge cells in the forefront vector when deciding the advancing direction. Our algorithm could potentially fail to capture the optimal path in the narrowed band, especially when the scores in the S vector are almost flat, which is likely to occur when the gap penalty is small or zero. We evaluated the effects of small gap penalties on the recall rate. Both the gap open penalty *G*_*o*_ and the gap extension penalty *G*_*e*_ varied independently from 0 to 5, with the match award and the mismatch penalty fixed to (*M*, *X*) = (2, 2). The results (Fig 4(c)) showed that setting the gap extension penalty to zero severely degraded the recall rate. A small but non-zero gap insertion penalty did not affect sensitivity when the gap extension penalty was greater than or equal to 2. This indicated that the algorithm also works well with the linear-gap-penalty function, where the gap penalty function is expressed in the form of *g*(*k*) = *k* · *G*, as well as with more general affine-gap-penalty functions. We also note that parameter combinations with non-perfect recalls are impractical, and will therefore not be used in real analyses. (1) *G*_*e*_ = 0 implies that we can insert nearly arbitrarily large gaps without incurring a significant penalty, and (2) (*G*_*o*_, *G*_*e*_) = (0, 1), (1, 1), (2, 1) all have a positive expected score for the alignment of two random sequences (four bases occurring equally and independently; Vingron and Waterman (1994)).

#### 3.1.3 Aligning longer sequences

Next, we evaluated the effect of the lengths of query sequences on the recall rate. If we use a statically banded DP, in which the band is determined before the values in cells are filled in, the band width required in order to capture the optimal path in the narrowed band grows by 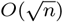 (where *n* is the expected number of insertion/deletion (indel) errors in a read, assuming that insertions and deletions occur at the same probability and independently). We hypothesized that there is a certain fixed-size band width that can efficiently capture the optimal path when the adaptive banded DP algorithm is used in empirical settings. Sets of simulated sequences of various lengths within the range of [100, 10000] were fed into our algorithm with three different band widths (*W* =16, *W* = 24, and *W* = 32). The results were then compared with those of the full semi-global alignment in order to determine whether they yielded comparable results. The mean sequence identity varied within the range of [0.6, 0.95], and the score parameters were set to (1, 2, 2, 1). The results (Fig 4(d)) show that the 16-cell-wide and 24-cell-wide bands may not be sufficient for perfectly aligning long (e.g., > 500) sequences with lower identity (~ 0.7). The 32-cell-wide band output the same result as the full semi-global alignment algorithm.

#### 3.1.4 Indel tolerance

Owing to the nature of banded DP algorithms, our algorithm is expected to fail to find the optimal score when a large indel appears in the true pairwise alignment. We evaluated the impact of indels in query sequences on the recall rate. We simulated an insertion by inserting a random sequence of a specific length into the reference-side sequences. Sequence pairs of a simulated PacBio read and its corresponding region in the reference genome were generated with the parameters *L* = 1, 000 and *I* = 0.75. Next, a random sequence of a specific length (denoted by *l*) within the range of [0, 100] was inserted into the reference-side sequence at a position within [100, 600] bp from its 5’-end. Considering that our algorithm runs basically symmetrically between the two sequences to be aligned, we tested only insertions; however, our results can likely be generalized to deletions as well. The score parameters (*M*, *X*, *G*_*o*_, *G*_*e*_) = (1, 2, 2, 1) were used. The results (Fig 4(d)) suggested that our algorithm can find the optimal score with a probability of nearly 100% when the length of the inserted gap, *l*, is less than *W* − 4.

### Performance on real datasets

We aimed to compare our algorithm with all existing extension alignment algorithms that are used in open-source long read aligners. Since our algorithm is an extension alignment algorithm, not a standalone aligner, we cannot directly compare our algorithm with existing standalone aligners. Instead, we extracted an extension algorithm part of existing aligners, and then compared them with our algorithm. First, we excluded some of long read aligners such as Kart (Lin and Hsu, 2017) and GraphMap (Sović *et al.*, 2016), because they do not extend long from an outermost seed in a chain so they do not contain an extension alignment algorithm. We also excluded BLASR (Chaisson and Tesler, 2012) and BWA-MEM (Li, 2013) (and minimap2 (Li, 2017)) because they extend a (outermost) seed (in a chain) always until the ends of query sequences and therefore they are not suited to long extension in local alignment. Note that it is an open question and beyond the scope of this paper whether the “pure seed-and-extend strategy” (BLAST+, BWA-MEM, etc.) is better than the “seed-and-chain-then-fill” strategy (BLASR, minimap2, etc.) in general. At least, we needed a long extension alignment algorithm for aligning reads from highly repetitive genomes, for which the latter is not likely to work well.

NCBI BLAST+ (Camacho *et al.*, 2009) and LAST (Kielbasa *et al.*, 2011) use largely the same algorithm for long extension, and therefore we extracted the extension algorithm from NCBI BLAST+ only because we did not observe an essential difference between their implementations. We call it “BLAST X-drop DP” hereafter. Because the BLAST X-drop DP algorithm solves exactly the same problem as our algorithm does, it is used as a baseline. We excluded NanoBLASTer (Amin *et al.*, 2016) for the following reasons: (1) it is designed for too noisy reads from old MinION sequencers; accepting 45% of sequencing errors seemed to put too much burden on the computation time. (2) its block-wise banded alignment algorithm is greedy so it does not try to compute an optimal path as the SWG algorithm does. (3) standalone NanoBLASTer fails with segmentation fault too often in our experiment, which made it difficult for us to work with its implementation.

Another algorithm from which we extracted an implementation is one of popular long read aligners, DALIGNER (Myers (2014)). DALIGNER is a long read aligner designed for finding alignment between long and noisy reads. Its core algorithm, the Myers’ wavefront algorithm (Myers, 1986), cannot be directly compared with our algorithm because the wavefront algorithm does not accept affine gap penalty; the wavefront algorithm can be considered as a special case of SWG-based extension algorithms. Nonetheless, we included the extension algorithm in DALIGNER (wavefront, hereafter) as a guide for aligner developers. In all, we compared the recall rate and the computation time of the adaptive banded DP with the BLAST X-drop DP algorithm and the wavefront algorithm.

#### Preparing input data

We selected as datasets whole-genome sequencing reads of a human genome (NA12878) from two single-molecule sequencers, PacBio RS II and Oxford Nanopore MinION (Zook *et al.*, 2014; Jain *et al.*, 2017). Since there is no way to have the ground truth for real datasets, we used the alignment results by BWA-MEM as ground truth because there are publicly available precomputed alignments, without which it may take months for preparation especially for the Nanopore reads. In each dataset, each alignment record by BWA-MEM was converted to a *read-reference sequence pair* that consists of the read sequence and the corresponding genomic region. We used a “virtual seed” setting, where a virtual seed is placed right next to the end of aligned sequences. To eliminate any bias of seeding algorithms or seeding positions, the read-reference sequence pairs were aligned by the four algorithms from randomly chosen end (3’- or 5’-end) toward the other end. To make the situation more realistic and include the effect of the X-drop termination in the evaluation, we added long random sequences to the tail (head when the alignment direction is opposite) of sequences so that the extension alignment always ends in the middle of a given sequence pair, not at the very end of the sequence pair. Since random sequences may have a small positive alignment score, this might change the optimal alignment a little bit but the impact is negligible. The MinION reads were obtained from the Nanopore WGS Consortium website. We only used reads aligned with Chr 20, which were the only reads basecalled by a newer algorithm called “Scrappie,” because those reads better reflect the latest error profile of MinION. According to the BAM header, the MinION reads were aligned with the GRCh38 reference genome (Schneider *et al.*, 2017) by BWA-MEM version 0.7.15-r1142-dirty with ONT 2D setting (“-xont2d”). The PacBio reads were obtained from the website of Genome in a Bottle Consortium. To make the alignment condition as close to the MinION reads as possible, the PacBio reads from Chr 20 were realigned by BWA-MEM with the PacBio setting (“-xpacbio”). Only primary alignments were used; secondary alignments were discarded. We randomly sampled 100, 000 sequence pairs from each dataset because it is a sufficient amount for evaluation.

#### Algorithm implementations and parameters

A simplified implementation was used for the BLAST X-drop DP algorithm since the implementation in NCBI BLAST+ contained unnecessary code for our experiment. We removed unnecessary code (e.g., code for protein alignment) and unused variables in the input and output data structures. The obtained source code is minimal so that we expect that it runs faster than or at least at the same speed to the original implementation. We also developed a SIMD-vectorized implementation of the BLAST X-drop DP based on our scalar implementation to observe the net contribution of the adaptive banded algorithm apart from the contribution of SIMD. The vectorized BLAST algorithm calculates the values of the cells in the DP matrix always in a vectorized manner, covering all the cells that are supposed to be calculated by the original BLAST+; in other words, it calculates broader area in the DP matrix than the scalar version. The way vectors are tiled was the same to one in Myers (1999). The implementation of the wavefront algorithm was extracted from the DALIGNER package (commit 84133cb). All the programs were compiled by Intel C Compiler 17.0.1 with an optimization flag “-O3.” The score parameters were set to (*M*, *X*, *G*_*o*_, *G*_*e*_) = (1,1, 1, 1) for the adaptive banded DP and the BLAST X-drop DP, following the ONT 2D and the PacBio setting of BWA-MEM used in the ground-truth alignments. The X-drop threshold was set to *X* = 70, which is equivalent to the default value (*X* = 100 bit) of NCBI BLAST+, for the adaptive banded and the X-drop DP algorithms. On X-drop threshold conversion, a parameter set (*M*, *X*, *G*_*i*_, *G*_*e*_) = (1, 1, 2,1) was used instead of the parameters used in the BWA-MEM mapping because it was not supported in the current version of NCBI BLAST+. The average correlation for the wavefront algorithm was set to 0.7, which is the default value of DALIGNER and also equals to the lower bound of the supported range (0.7 to 1) of the algorithm.

#### 3.2.1 Recall benchmark

We measured the recall rates by the four algorithms. The recall rate is defined as the number of sequence pairs for which an alignment of the same score or higher was found. The band width was varied between 32 and 256 for the adaptive banded DP algorithm. The adaptive banded DP hit a better balance between speed and sensitivity over the other algorithms on the both datasets; a better recall rate was achieved with the equivalent computation time and faster calculation was achieved with the equivalent recall rate (Fig 5). The same results are shown in Table 1 for three band widths for the adaptive banded DP, *W* = 64, 96, and 128 (dotted on lines in Fig 5(a) and (b)). The wavefront algorithm was the fastest among all, while the recall rates were low compared to the others.

**Fig. 5.**
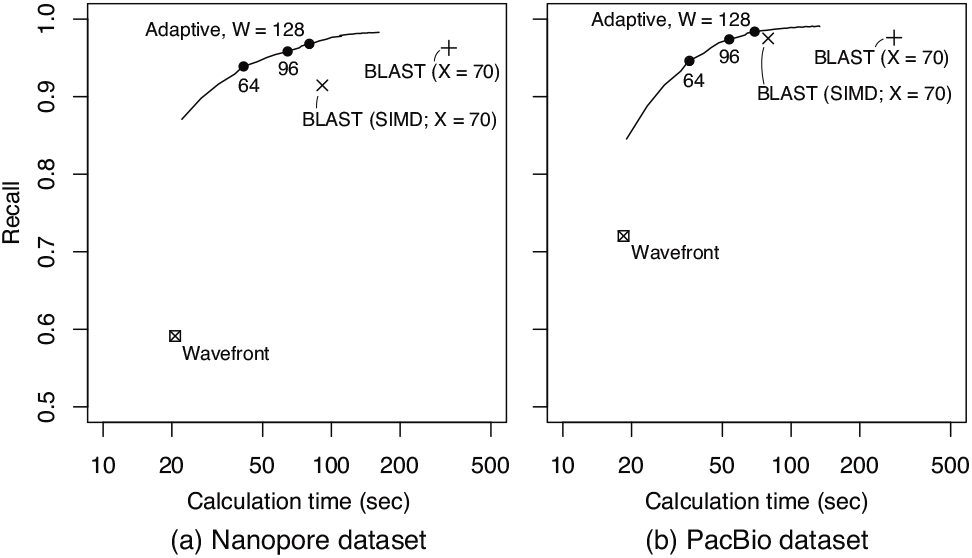
The recall rates for the three algorithms. The lines show the trend of the recall rates with respect to the calculation time by the adaptive banded DP with *W* = 32 to 256 (step size: 8); points are shown only for *W* = 64, 96,128.

**Table 1.** The recall rates and the computation time for the four algorithms.

|  |  | Nanopore |  | PacBio |  |
| --- | --- | --- | --- | --- | --- |
|  |  | Recall (%) | Run time (s) | Recall (%) | Run time (s) |
| Adaptive | W = 64 | 93.85 | 40.8 | 94.58 | 35.5 |
|  | W = 96 | 95.78 | 63.6 | 97.34 | 53.0 |
|  | W = 128 | 96.77 | 79.2 | 98.34 | 68.5 |
| BLAST | X = 70 | 96.29 | 327.2 | 97.61 | 282.6 |
| BLAST (SIMD) | X = 70 | 91.46 | 91.3 | 97.51 | 79.3 |
| Wavefront |  | 59.13 | 20.7 | 72.03 | 18.4 |

#### 3.2.2 Speed benchmark

We evaluated the matrix calculation performance of several algorithms with respect to different query lengths using the same implementations. The band width was fixed to 96 for the adaptive banded DP so that the recall rate becomes similar to one by the X-drop DP with *X* = 70. In addition to the four implementations, the Farrar’s SIMD-vectorized SWG algorithm (Farrar, 2007) was added to the comparison targets as a guide for aligner developers. Although it is a non-banded, full-sized DP algorithm, it is adopted in commonly used short-read alignment tools (BWA (Li and Durbin, 2009) and Bowtie 2 (Langmead and Salzberg, 2012)) as a fast SWG algorithm. The SSE4.1 implementation of the semi-global variant of the Farrar’s algorithm was obtained from Parasail library (commit 3d8b4ee; Daily (2016)) because it is highly optimized for various instruction sets.

The query sequence length *L* was varied from 1 bp to 10 kbp, by trimming the tail of the 10 kbp or longer sequence pairs that were randomly sampled from each dataset. The results, shown in Figure 6(a) and (b), were quite similar between the two samples. The 96-cell-wide adaptive banded DP was consistently faster than the scalar and the SIMD-vectorized BLAST X-drop DP algorithms by 5.1 and 1.5 times, respectively. The Farrar’s algorithm was the fastest for short sequences (roughly *L* < 200), while it was the slowest for long sequences (*L* > 2, 000) due to its *O*(*L*^2^) complexity. The wavefront algorithm was the fastest for long sequences (roughly *L* > 2, 000).

**Fig. 6.**
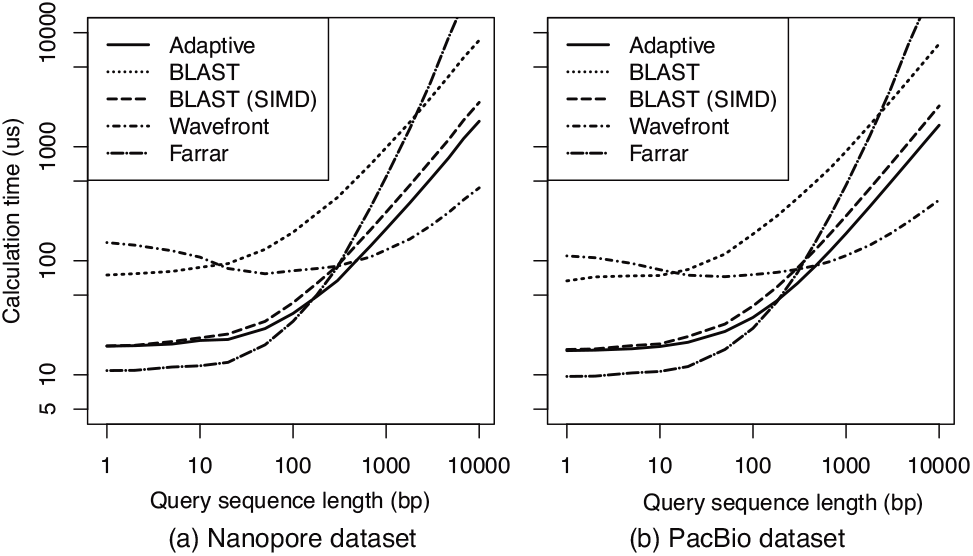
The computation time for matrix fill per alignment and the recall rates at 10 kbp for five algorithms for the PacBio (a) and Nanopore (b) samples: the BLAST X-drop DP (BLAST) and the SIMD-vectorized BLAST X-drop DP (BLAST-SIMD) with *X* = 70, the and 96-cell-wide SIMD-vectorized adaptive banded DP (Adaptive), the Myers’ wavefront algorithm in DALIGNER (Wavefront), and the Farrar’s SIMD-vectorized in Parasail library (Farrar). The times at 1 bp can be largely considered as the time needed for finding the end of an alignment.

## 4 Discussions

We proposed an adaptive banded DP algorithm for nucleotide alignment that calculates only the cells in a dynamically determined band of constant width in the DP matrix. Even though the new algorithm uses bands of constant width regardless of sequence lengths, it retains a high sensitivity in long extension, suggesting that the new algorithm is immediately useful for resequencing analysis and de novo assembly. The new algorithm runs fast because it is friendly to SIMD vectorization because the band width can be always a multiple of the SIMD width and because anti-diagonally placed vectors eliminated dependences between elements in a single vector. Other algorithms, including our vectorized BLAST X-drop off algorithm and the Rognes’s algorithm (Rognes and Seeberg, 2000), have dependences between elements in a vector, for which serial calculation of cell values is needed. The difference will increase, as newer processor with increased vector width (e.g., one with 512-bit-wide Advanced Vector eXtension instructions) are released. The use of an SIMD shuffle instruction for generating the score profile vector allowed us to execute the operation in a vectorized form, although it cannot be directly extended for protein alignment because the score matrix for amino acids does not fit in a single register.

It should also be noted that the 16-bit-wide implementation of our algorithm, with a minor modification, is sufficient for supporting (virtually) infinitely long query sequences. That is, the actual scores in each vector can be represented by a pair of a potentially large base values of 64-bits and an offset value of 16-bits from the base value, such that the elements (i.e., offset values) in vectors fit within the 16-bit range. We can thereby ensure that the range of scores in each vector can be bounded within a moderately small range, assuming that the absolute values of score parameters are small integers.

Finally, we would like to point out that it is possible to port our implementation to other architectures because we designed it with portability in mind. Power and AArch64 architectures have their own SIMD instruction sets, AltiVec and NEON, both of which have the basic arithmetic and 16-element shuffle operations required for our algorithm. As for the current GPU architectures (e.g. NVIDIA), it is expected that our constant-wide adaptive banded DP algorithm is more suited than other existing semi-global alignment algorithms, where each SIMD lane can be statically assigned to synchronous threads.

## 5 Availability of the code

The implementation of the algorithm and the benchmarking scripts are available at https://github.com/ocxtal/adaptivebandbench.

## 6 Acknowledgements

This work was supported in part by MEXT KAKENHI Grant Number 16H06279. We also thank Professor Shinichi Morishita for his kind and insightful comment on this manuscript.

## Footnotes

1 Strictly speaking, we have two center cells when W is odd, but here we define the center as either of them.

